# The *FIBRILLIN* multigene family in tomato, their roles in plastoglobuli structure and metabolism

**DOI:** 10.1101/2023.01.05.522848

**Authors:** Juliana Almeida, Laura Perez-Fons, Margit Drapal, Kit Liew, Paul D. Fraser

## Abstract

Plastoglobuli (PG) are plant lipoprotein compartments, present in plastid organelles. They are involved in the formation and/or storage of lipophilic metabolites. FIBRILLINs (FBN) are one of the main PG-associated proteins and are particularly abundant in carotenoid-enriched chromoplasts found in ripe fruits and flowers. To address the contribution of different FBNs to isoprenoid sequestration and PG function, a multiplex gene editing approach was undertaken. Analysis of single and high-order *fbn* mutants for the major PG-related FBNs in tomato, namely *SlFBN*1, *SlFBN*2a, *SlFBN*4, *SlFBN*7a, revealed functional redundancy. High order *fbn* mutants displayed phenotypes associated with abnormal isoprenoid accumulation, and aberrant PG formation and morphology. Lipidomic analysis highlighted broader changes in lipid metabolism. Paralog-specific roles were also observed and included the regulation of specific isoprenoids (e.g., plastochromanol) and control of plastidial esterification capability by SlFBN7a. Collectively, the results support both structural and regulatory roles of SlFBNs in PGs. Our findings expose fundamental aspects of metabolic compartmentalisation in plant cells and the importance of lipoprotein particles for their plastid metabolism/physiology.

**Significance statement:** In the chromoplast of ripe tomato fruit and flower, plastoglobuli (PGs) are associated with several important biotechnological traits, due to their functional involvement in metabolism, developmental transitions, and environmental adaption. FIBRILLINS (FBN) are a multigene family of proteins that are collectively major components of the PG. Using a multiplex CRISPR-Cas9 approach single and high-order *fbn* mutants have been developed. Functional redundancy amongst the members of the *FBN* multigene family was evident, but also paralog specific functions/influence. Aberrant plastoglobuli formation and altered lipid metabolism are evident among *fbn* mutants. Characterisation of this resource has shed light on the functional role of FBN and their role in PG formation. This strategy offers new potential for the development of nutritional enhanced and climate resilient crops.

## INTRODUCTION

The role of lipid droplets and lipoproteins in maintaining lipid homeostasis has been studied extensively in animal systems due to their involvement in cardiovascular diseases and obesity (1). Similar lipoprotein structures are also present in the plastids of plants. However, their function remains comparatively poorly understood (2). Plant plastids, including photosynthetically active chloroplasts and other types, contain dynamic lipid storage sub-compartments termed *plastoglobuli* (PGs). PG number, size and composition vary depending on plastid differentiation and type being linked to plant developmental transitions. PG characteristics can also change as a result of stress responses (3)(4). Recent evidence has supported that the PG is not only a sink for lipids but an active metabolic hub containing key metabolites of biotechnological importance (5, 6).

Structurally, PGs are surrounded by a membrane lipid monolayer, encapsulating a neutral lipid core. Triacylglycerides (TAGs) can predominate in the core, depending on plastid type, but also present are isoprenoids (carotenoids, tocopherols and other prenylquinones) (7). A specific set of proteins coat the PG surface which also greatly vary depending on cell type (3). A major structural protein class found in PGs are FIBRILLINs (FBN). These plastid-specific proteins are encoded by a multigene family whose members contain a typical plastid lipid-associated protein (PAP) domain. They do, however, lack integral transmembrane domains but possess amphipathic helices predicted to bind to PG lipid monolayer (8, 9). FBNs also contain a conserved lipocalin-like motif which has been suggested either to participate in the binding and transport of small hydrophobic molecules (8, 10).

In the model flowering plant *Arabidopsis*, 14 FBNs have been identified; seven of which are considered PG-associated proteins, while the others are located primarily in thylakoid membranes or stroma of chloroplasts (3, 11). Functionally, FBNs are involved in different plant processes ranging from photosynthesis to colour acquisition during organ development and response to stress conditions (12–14). Moreover, FBNs can fulfil key structural roles associated with metabolic pathways (10, 15).

FBNs are best known for their role in chromoplasts, the coloured type of plastids. Typically occurring in flowers and fruits, chromoplasts remodels their metabolism to synthesise and store large amounts of carotenoids, isoprenoid-derived compounds with high nutritional and industrial value (16, 17). FBN1/PAP and its homologs in different species are required for the formation of carotenoid-sequestering structures occurring during fruit ripening as PGs and fibrils (13, 18–21). Enhanced-carotenoid plant genotypes produced by biotechnological interventions typically exhibited altered chromoplast structure with increased PG number, highlighting the metabolic composition influence on plastid morphology (22–25).

Despite multiple studies implementing FBNs with PG formation and dynamics, the underlying mechanisms of how FBNs fulfil this role remain poorly understood. It has been proposed that PG-targeted FBNs may control globule size, preventing PG coalescence, and favouring their clustering (26). Given the multigene feature of FBNs, questions remain as to which FBN members act on PG formation, and/or which affect PG core lipid remodelling, leading to changes in plant metabolism. Indeed, individual FBN contributions is challenging to interpret because several members occur simultaneously in plant cells.

In the present study, the independent and complementary roles of the major PG-related FBNs have been investigated by adopting a multiplexing gene editing approach in tomato. Metabolomic, proteomic and cellular ultrastructural analysis has provided a deeper insight into the functional divergence of the *FBN* gene family and how they can be exploited in future biotechnological applications.

## RESULTS

### Putative plastoglobular FBNs associated with tomato chromoplast development

Using the *Arabidopsis thaliana* FBN (AtFBN) protein sequences (3), 15 tomato *FBN* (*SlFBN*) genes, were identified in total covering all AtFBN groups. For comparison, homologous FBN sequences from pepper (*Capsicum annuum, CaFBN*s), another solanaceae species with fruits highly specialized for carotenoid storage, were also retrieved. *SlFBN* orthologs of *AtFBN*s encoding PG-targeted proteins (11), namely *SlFBN1/CHRC, −2a, −2b, −4, −7a, −7b, −8* were determined by phylogenetic analysis. Notably, paralogs pairs primarily clustered within each plant family, supporting lineage-specific expansions (Fig. 1A).

**Fig. 1:**
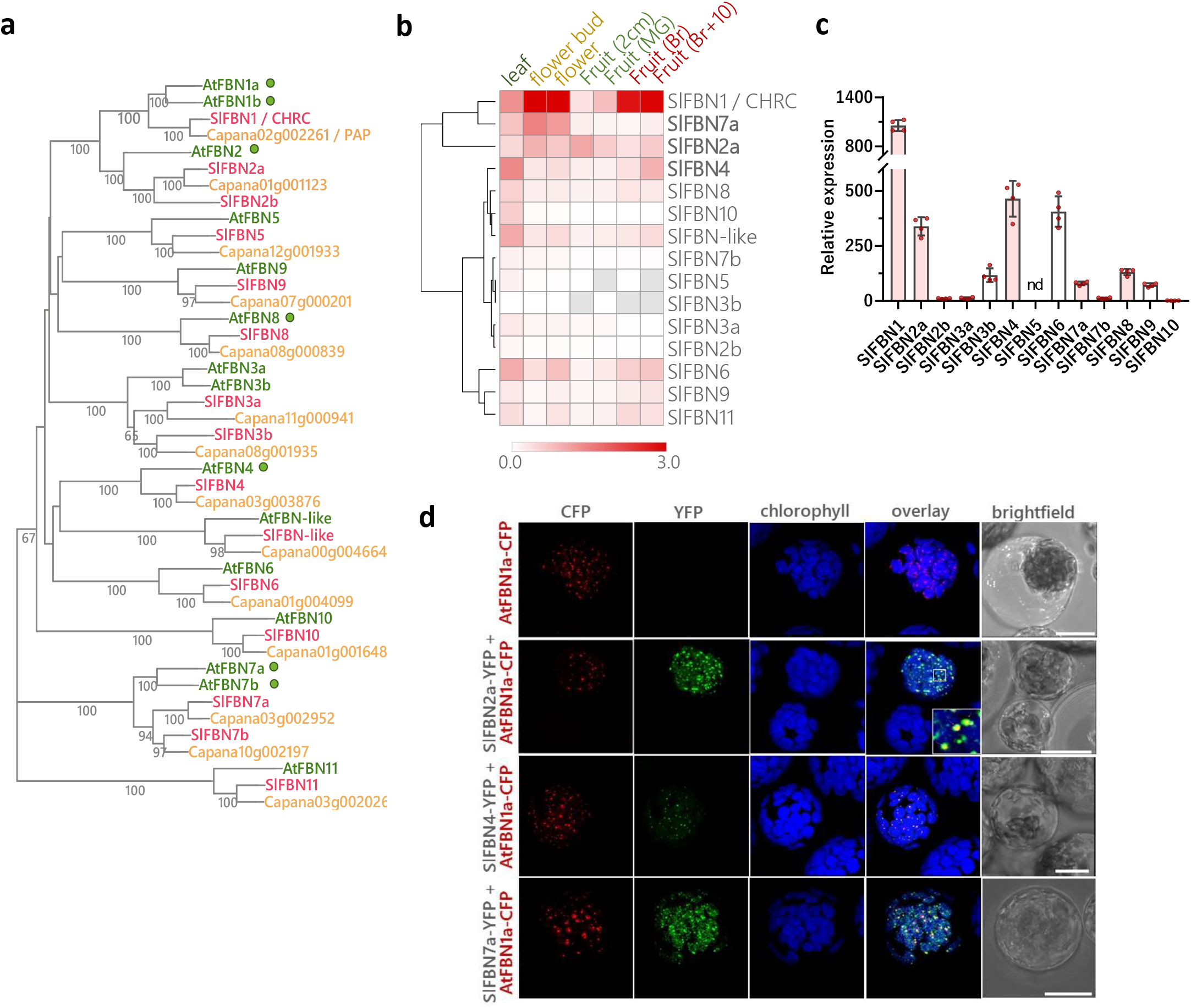
Identification tomato FIBRILLINs (SlFBNs), expression profile and subcellular localisation.

Of the seven PG-related *SlFBN*s transcripts levels of five members (*SlFBN1, −2a, −7a, −4*, −*8*) were highest at carotenoid-enriched stages of flower and fruit, based on transcriptome database queries (Fig. 1B). Their expression profile was further confirmed by qPCR analysis in fruit (Fig. 1C). *SlFBN1/CHRC* was confirmed as the most highly chromoplast-expressed *SlFBN* (19), and correlated to *SlFBN2a* and −*7a* expression (Fig. 1B). Besides, *SlFBN4* transcripts also accumulated as tomato fruit ripening advanced.

Pepper transcriptomic data (27) was also interrogated to check *CaFBN* transcripts abundance across the fruit development. Interestingly, PG-related Ca*FBN* homologs had expression pattern similar to tomato (highly expressed at chromoplast-enriched tissues/stages) (SI Appendix, Fig. S1). *CaFBN1/PAP* was the predominant Ca*FBN* expressed in fruits. Despite differences in flowers due to lack of chromoplasts in pepper petals, high expression rates of *CaFBN4*, *CaFBN2* and *CaFBN7a* are detected at the late stages of pepper fruit ripening, arguing for their role in carotenoid sequestration.

Based on the above orthologs and expression profiles, *SlFBN2a, SlFBN4* and *SlFBN7a* emerged as candidates involved in PG biogenesis and carotenoid sequestration, potentially complementing SlFBN1/CHRC function in chromoplasts. To first check their subcellular localisation, a yellow fluorescent protein-tagged of each SlFBN candidate (SlFBN-YFP) was transiently co-expressed with the plastoglobular marker AtFBN1a (28) in protoplasts. The signal fluorescence of all SlFBN-YFP fusions was restricted to the plastids. While SlFBN4-YFP fluorescence greatly co-localised with PG marker, the patterns of SlFBN2a-YFP and SlFBN7a-YFP suggested a more diffuse distribution of these proteins within the plastid not restricted to PG (Fig. 1D). Together, the results suggest that SlFBN2a, −4, −7a can be associated with PGs, and likely have a role in PG formation and composition.

### Generation of single and high-order *fbn* mutants by CRISPR-Cas

To address the contribution of *SlFBN*2a, −4, −7a in PG formation and isoprenoid sequestration, single loss-of-function mutants for the candidate *SlFBN*s were generated via CRISPR-Cas technology (Fig. 2, SI Appendix, Table S1). To avoid functional redundancy within the FBN family, a multiplexing approach was also undertaken to simultaneously deliver combinations of mutant alleles for *SlFBN*1, −2a, −4 (triple mutants) and *SlFBN*1, −2a, −4, −7a (quadruple mutants) (SI Appendix, Fig. S2).

**Fig. 2:**
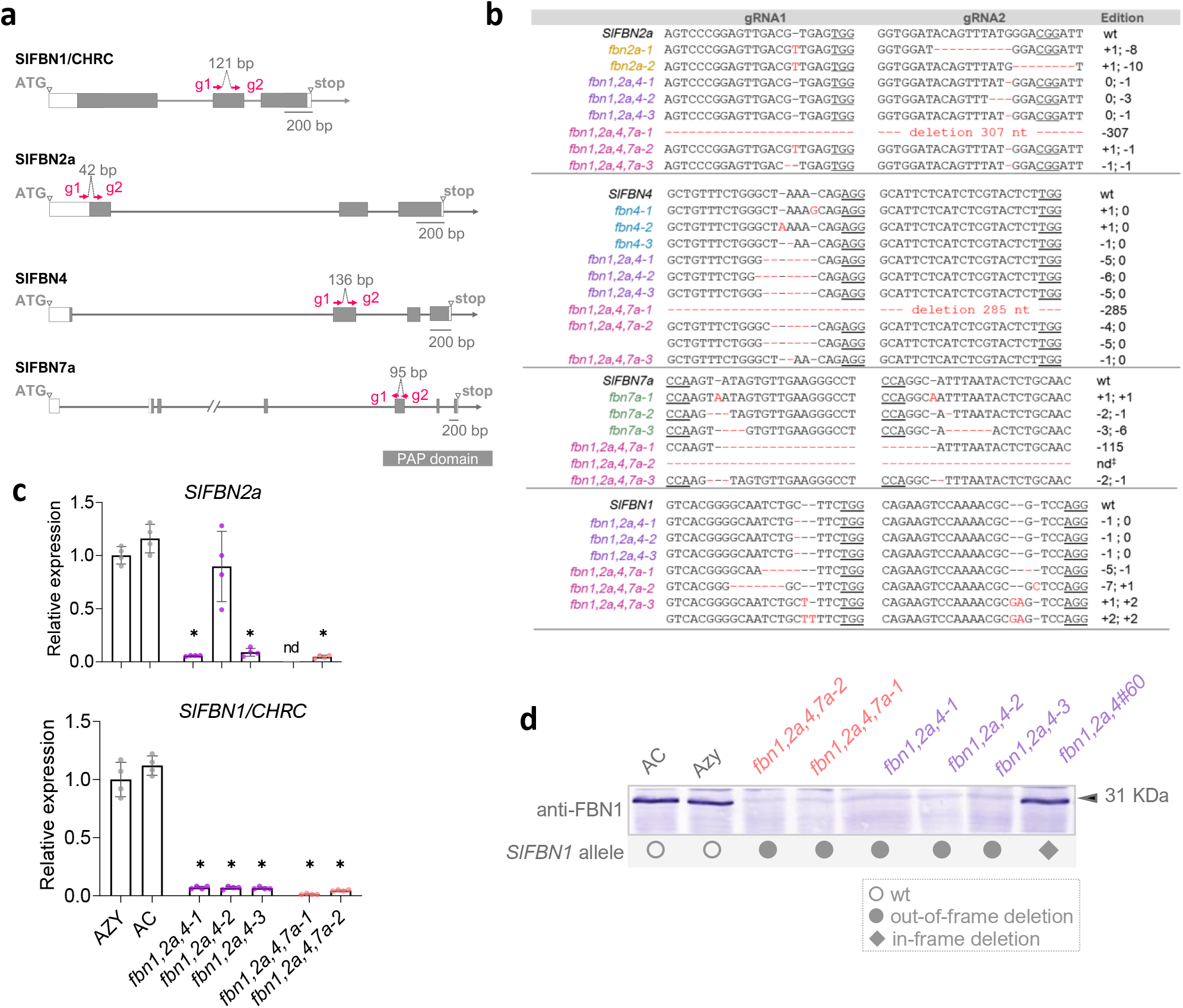
Gene-edited *fbn* mutants generated by CRISPR-Cas.

Sequencing of the targeted genomic regions confirmed edited alleles in primary T_0_ transformants obtained through single and multiplexing strategies. Editing efficiency ranged from 67 to 100% (SI Appendix, Table S2). Mutations occurred in different combinations, chimera, biallelic and homozygous, being mostly short indels upstream PAM sites. Large out-of-frame deletions (> 100 bp) were also detected at a lower frequency. Edited alleles harbouring frameshift or nonsense (premature stop codon, PTC) mutation were predicted as knock-outs. Lines carrying in-frame deletion alleles, which could either retain functionality or be loss-of-function, were also selected for comparison when available. Possible off-target sites for the gRNAs examined in T_0_ and T_1_ plants (SI Appendix, Table S1) were undetected, supporting the high specificity of the gRNAs designed. T_2_ Cas-free homozygous/biallelic *SlFBN* plants were successfully established and used for further comprehensive phenotyping and molecular characterisation (Fig. 2 and SI Appendix, Table S3). Since transcripts with PTC introduced by gene editing can be targeted by nonsense-mediated decay pathways, by which nonsense mRNAs were destroyed (29), expression of *SlFBN*1/*CHRC* and *SlFBN*2a, both highly expressed in fruits, were checked by qPCR. Levels of PTC-containing mutant transcripts were reduced in *fbn* fruits compared to theirs control (Fig. 2C). Additionally, FBN1 protein levels were checked by immunoblotting using an antibody to CaFBN1/PAP (21). *SlFBN*1 was exclusively detected in fruits from edited lines carrying either wild-type or in-frame deletion alleles (Fig. 2D).

Visual inspection of *fbn* mutants revealed no obvious difference in fruit characteristic red colour of *fbn* mutants but altered flower pigmentation. The typical bright yellow colour failed to develop particularly in *fbn* mutants harbouring *SlFBN7a* defective alleles (single *fbn7a* and quadruple *fbn1,2a,4,7a*) (Fig. 5). Triple and quadruple mutants, under non-stressed growth conditions, had some altered plant morphological traits. These morphological changes consisted of a shorter internode length compared to their controls and visually leaf areas were reduced in the quadruple mutants (SI Appendix, Fig. S3). Small decreases in maximum quantum efficiency of PSII (Fv/Fm) were found in one line of quadruple *fbn* mutants.

### *SlFBN* homologs influence the accumulation of carotenoids and other isoprenoids in plastids

To assess how *SlFBN* mutations influences plastidial isoprenoid metabolism, fruit and leaf isoprenoid profiles, were firstly examined by UPLC-PDA (Fig. 3 and SI Appendix, Table S4). Three fruit stages were analysed: mature green (MG), red ripe (B7) and late red ripe (B10). Carotenoid changes occurred predominantly in ripe fruits from high-order *fbn* mutants (Fig. 3B; SI Appendix, Table S4). The early carotene intermediates (phytoene, phytofluene and ζ-carotene) were mostly affected and reduced by 60% in the mutants compared to control (Azygous, Azy) at B10 stage. Unexpectedly, the decrease in lycopene, the main carotenoid accumulating in red fruit, was less pronounced (levels of 80% retained). B-carotene levels, however, were unaffected despite lower levels of its immediate precursor γ-carotene. In single *fbn* fruits, carotenoid content remained mostly similar to control at all fruit stages.

**Fig. 3.**
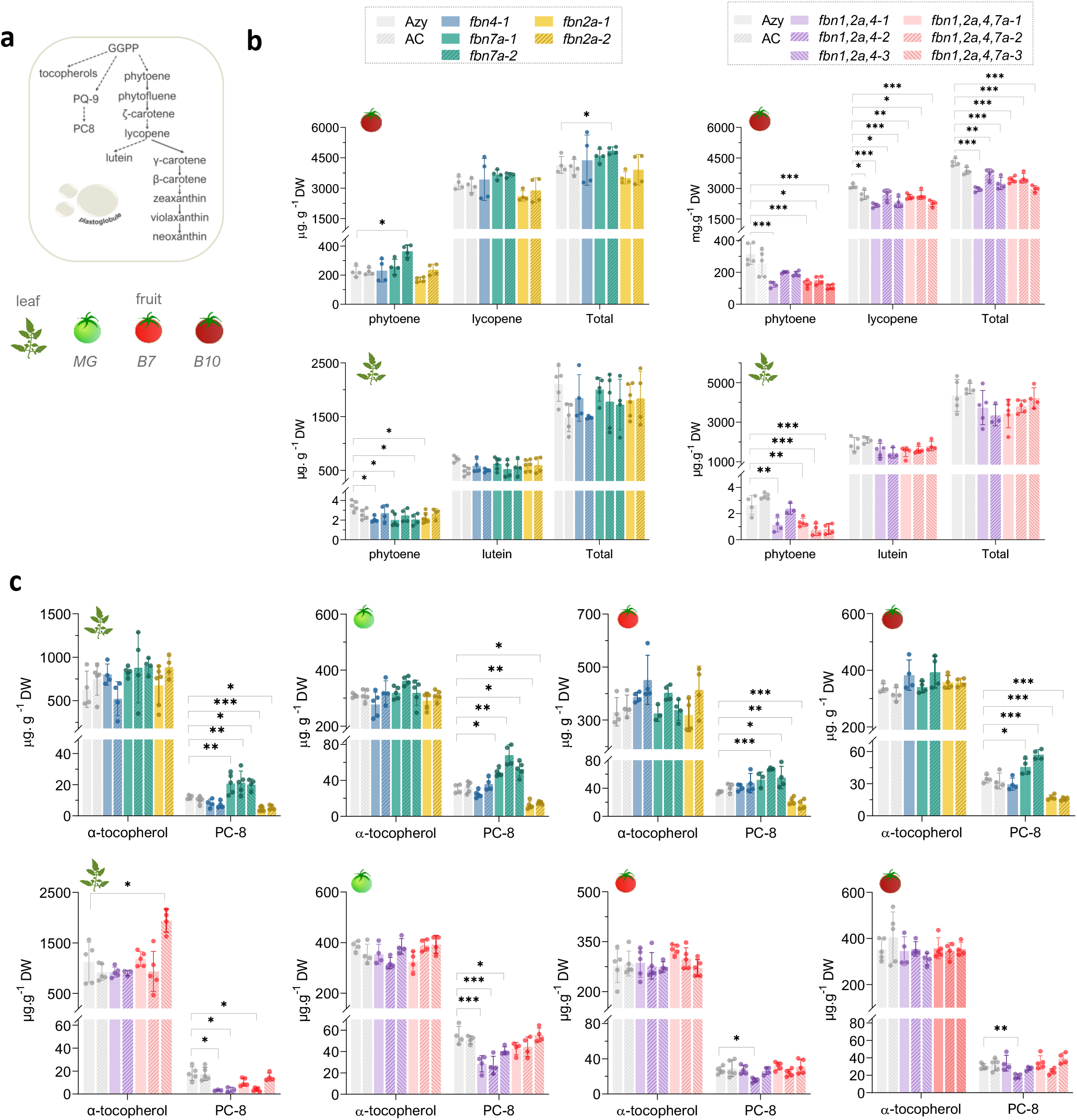
SlFBNs altered isoprenoid profile in tomato fruits and leaves of *fbn* mutants.

Other isoprenoids known to be sequestered in the PG core, such as tocopherol and quinone derivatives, responded differently to single or multiple losses of *SlFBN*s (Fig. 3B; SI Appendix, Table S4). Despite similar levels of α-tocopherol and plastoquinone (PQ-9), plastochromanol-8 (PC-8) significantly changed in *fbn* mutants. PC-8 is a PQ-9 derivative predominantly found at PG associated with stress response (30). While the lack of *SlFBN2a* approximately halved PC-8 levels compared to the control, deficiency in SlFBN7a had an opposite effect, favouring PC-8 accumulation with up to 2-fold increase. Of note, similar PC-8 levels observed among *fbn7a* lines carrying either in-frame (*fbn7a-3*) or nonsense alleles (*fbn7a-1, fbn7a-2*) suggested that all *SlFBN7a* editions likely derived *SlFBN7a* null alleles. In high-order *fbn* mutants, *SlFBN2a* and *SlFBN7a* defective alleles displayed compensatory interaction to modulate PC-8 levels though outcomes were specific to the organ and developmental stages. In triple mutants, PC8 levels were decreased in chloroplast-containing tissues, MG fruits (50%) and leaves (20%), compared to control, whilst no consistent effect was found in red fruits. In quadruple mutants, the addition of a defective *SlFBN7a* restored PC-8 levels in fruits at all stages. Together, these findings define mutation-specific effects of *SlFBN*s on levels of prenylquinones. While *SlFBN2a* favours PC-8 accumulation, *SlFBN7a* prevents it. To further substantiate these findings, an extra set of T_1_ *SlFBN*-edited lines harbouring Cas9-gRNA transgene (*Cas^+^*) with the following genotypes: triple (*Cas^+^/fbn1,2a,4*), quadruple (*Cas^+^/fbn1,2a,4,7a*) and double (*Cas^+^/fbn2a,4*) mutant (SI Appendix, Table S3). This additional set of *fbn* mutants allowed us not only to confirm the fruit profile found T_2_ Cas-free progeny but also to evaluate SlFBN1-specific contribution to isoprenoid profile by comparing *Cas^+^/fbn2a,4* and *Cas^+^/fbn1,2a,4* (SI Appendix, Table S4). PC-8 levels were similar between double and triple mutants, implicating SlFBN2a as the major positive regulator of PC-8 levels.

### Changes in isoprenoid sequestration in fruit chromoplasts of *fbn* mutants

*SlFBN*-dependent accumulation of isoprenoids may be linked to how these compounds are preferentially sequestered across plastid sub-compartments (24). To test this hypothesis, sub-plastidial chromoplast fractionations were isolated and analysed for isoprenoid contents. A differential sequestration pattern was observed in high-order *fbn* mutants. For carotenoids, the proportion of phytoene and phytofluene were decreased in the PG fraction of *fbn* mutants compared to control. Instead, these carotenes preferentially accumulated at fractions corresponding to membranes and crystals. Altered deposition of α-tocopherol and PC-8 across subchromoplast compartments was also observed (Fig. 4b). Accordingly, the relative composition PG fractions greatly changed compared to control when multiple *SlFBN* were perturbed (Fig. 4c). Together, these results suggest a reduced capacity of isoprenoid sequestration at PG caused by multiple *SlFBN* disruptions.

**Fig 4.**
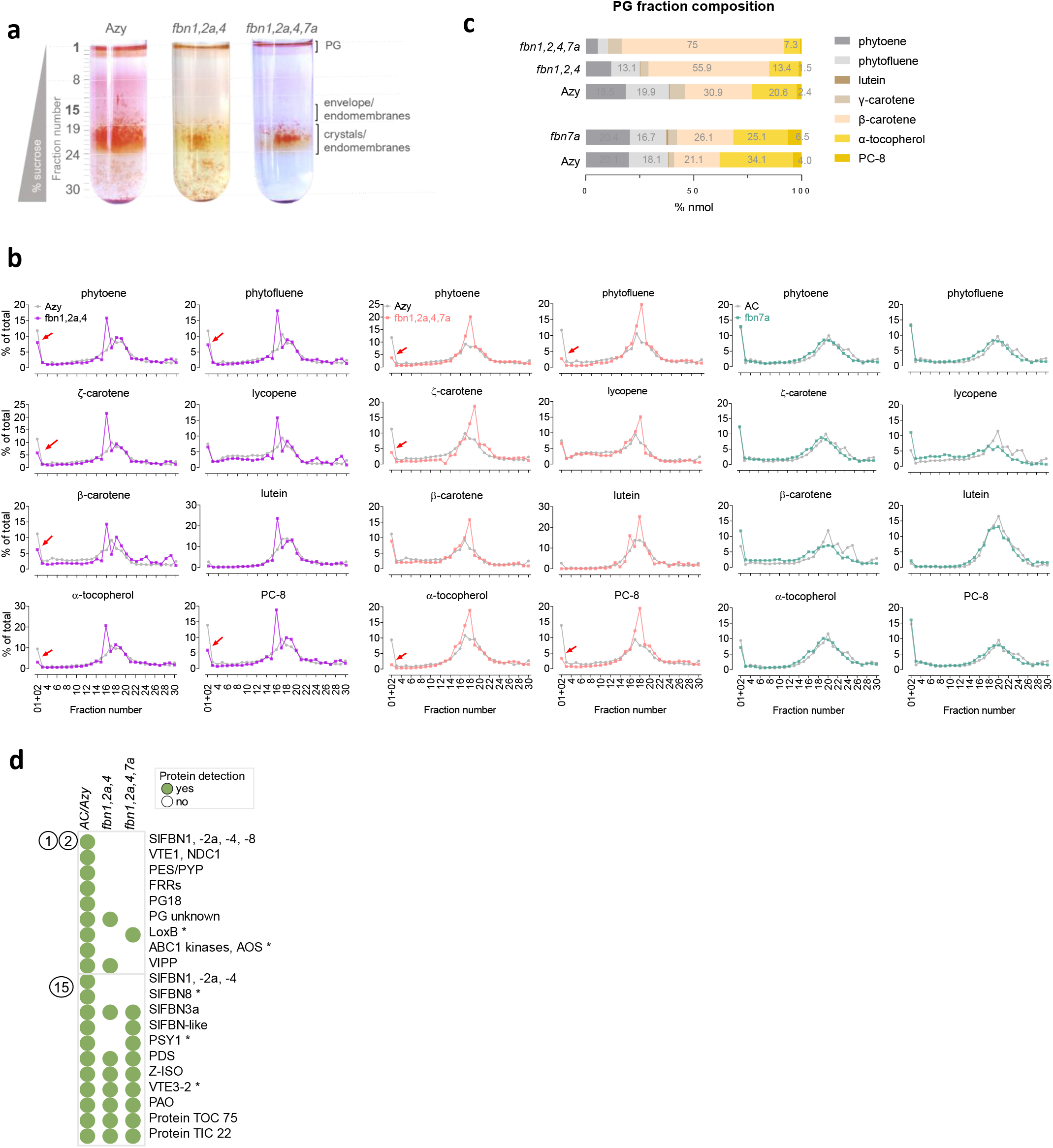
Sequestration of isoprenoids into subchromoplast compartments in *fbn* mutants.

### SlFBN7a modulates esterification capability in flower chromoplasts

The yellow pigmentation of tomato flowers is determined by xanthophylls in their esterified form esterified to medium-length fatty acids (C14:0-C16:0), forming mono- and di-esters (31, 32). While *fbn2a* and *fbn4* were typically bright yellow, *fbn7a* petals appeared pale (Fig. 5a). Quadruple mutants also failed to develop yellow pigmentation showing a more severe phenotype than *fbn7a*. Triple *fbn* mutant flowers only exhibited an apparent less intense yellow colour. To better understand the biochemical basis of the pale phenotype, detailed compositional analysis of carotenoids in petal extracts were performed using high-performance liquid chromatography (HPLC-PDA) to resolve carotenoids in free, mono- and di-ester forms (Fig. 5A, 5B; SI Appendix, Table S5). Carotenoid level and composition greatly changed in *fbn7a* petals. The total levels were decreased by ~50% relative to control. The free carotenoids became predominant in *fbn7a* compositional profile (3-fold increase), followed by a decrease in the proportion of mono- and di-esters (~70% and 65% of the values found in control, respectively). No significant changes in content nor the composition of *fbn2a* and *fbn4* petals were detected. Analysis of high-order *fbn* mutants further confirmed the major role of *SlFBN7a* in modulating the esterification capability of flower chromoplasts. Pale yellow petals of *fbn1,2a,4,7a* were like *fbn7a* and primarily accumulated free xanthophylls (> 4-fold higher versus control), while having lower levels of esterified xanthophylls. The decrease in mono- and di-esters proportions (45% and 30% of the values found in control) exceeded those observed in *fbn7a* solely (SI Appendix, Table S5), suggesting further disruption in the esterification capability when multiple SlFBNs are defective. Importantly, *fbn1,2a,4,7a* flowers had noticeable β-carotene quantities, which is often analytically masked by xanthophyll esters in non-saponified extracts (33). Instead, triple mutants, harbouring a FBN7a protein, maintained carotenoid composition, consistent with the FBN7a role in the regulation of esterification activity in flower chromoplasts.

**Fig. 5.**
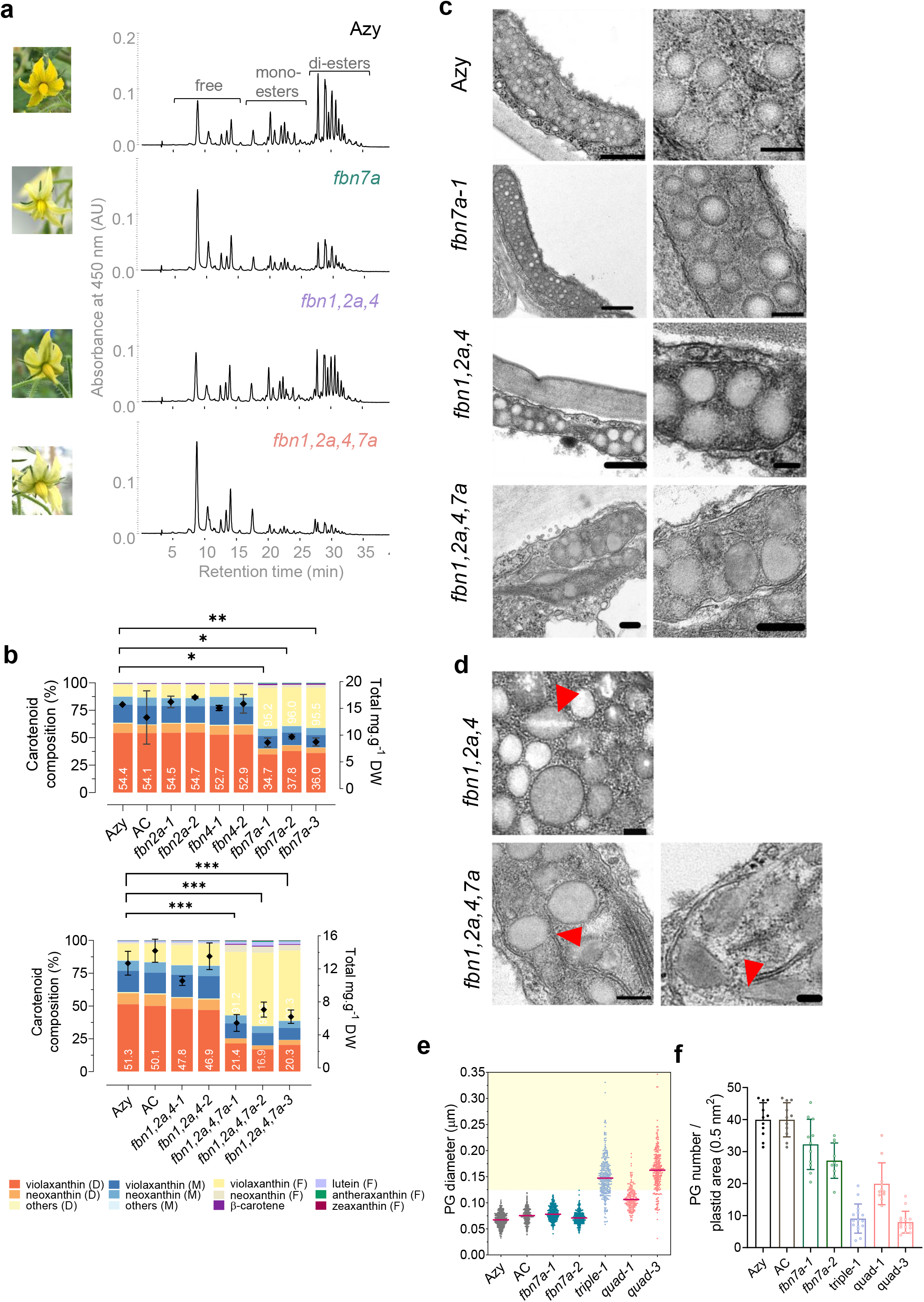
SlFBN7a regulates carotenoid esterification driven colour retention in tomato flowers, while SlFBN1, −2a, 4 affect PG morphology.

A similar compositional profile was also confirmed in the extra T_1_ *fbn* mutants (SI Appendix, Fig. S4, Table S5), although in this case, total carotenoid content of triple (*Cas^+^/fbn1,2a,4*) was significantly reduced compared to wild type (AC), which may reflect the absence of azygous comparators at this stage. Furthermore, total carotenoid amounts found in triple (*Cas^+^/fbn1,2a,4*) and quadruple (*Cas^+^/fbn1,2a,4,7a*) were equivalent, indeed lower than those found in double (*Cas^+^/fbn2a,4*) mutant, thereby confirming SlFBN1/ChrC as one of the main contributors for xanthophyll sequestration in flower chromoplasts.

### FBNs are required for PG formation and chromoplast development

To investigate the effect of PG-targeted SlFBN deficiency on PG formation and chromoplast architecture, petals and fruit pericarps were examined under transmission electron microscopy (TEM). The plastids in petals were present in different forms (31), ranging from chloroplast-like structure (early stage) to the fully re-differentiated carotenoid-rich chromoplasts. Early-stage plastids had thylakoids organised in grana and low numbers of electron dense PGs (Fig. 5C). Mature chromoplasts instead displayed the typical abundant small PGs (diameter ~60 nm), with a less electron-dense core where carotenoids are sequestered. In pale yellow *fbn7a* petals, PGs were similar to the control in size and morphology but appeared less in number (Fig. 5C-5F). High-order *fbn* mutants had chromoplasts with great variation in sub-structures, including thylakoid remnants, with few and supersized PGs (diameter > 140 nm, Fig. 5e). In some cases, their abnormal PGs seemed to be associated with the persistent dismantled thylakoids and showed a discontinuous lipid monolayer instead of a perfectly smooth surface. Additionally, they displayed crystalloid-like inclusions (*e.g*., lightly stained zones) in the form of filaments (Fig. 5D). Particularly in quadruple mutants, with predominantly non-esterified xanthophylls, PGs even assumed elongated to spindle-shaped forms, instead of round structures, filled with homogenous electron-dense filaments. Interestingly, early-stage plastids of high-order *fbn* mutants exhibited a typical chloroplast-like structure with only a few supersized PGs (SI Appendix, Fig. S5).

Unlike flowers, ripe fruits in tomato contain more diverse carotenoid-sequestering structures including endomembranes, carotenoid crystals and bigger PGs (34). The chromoplasts found in control samples displayed crystals of lycopene and β-carotene and several high-electron dense PGs ~ 100 nm in diameter. PGs of single *fbn2a* and *fbn7a* mutants changed in number and morphology, whereas *fbn4* chromoplasts remained similar to control (Fig. 6). In *fbn2a* mutants, supersized PGs (diameter > 200 nm) were observed at high frequency, with some of them displaying electron-lucent areas at their core. In *fbn7a* mutants, fruit PGs were found with a less electron-dense core and only a small proportion corresponded to supersized PGs (at least 25%). The changes in PG electron density are likely to be due to altered chemical composition, affecting extraction during preparation (35).

**Fig 6.**
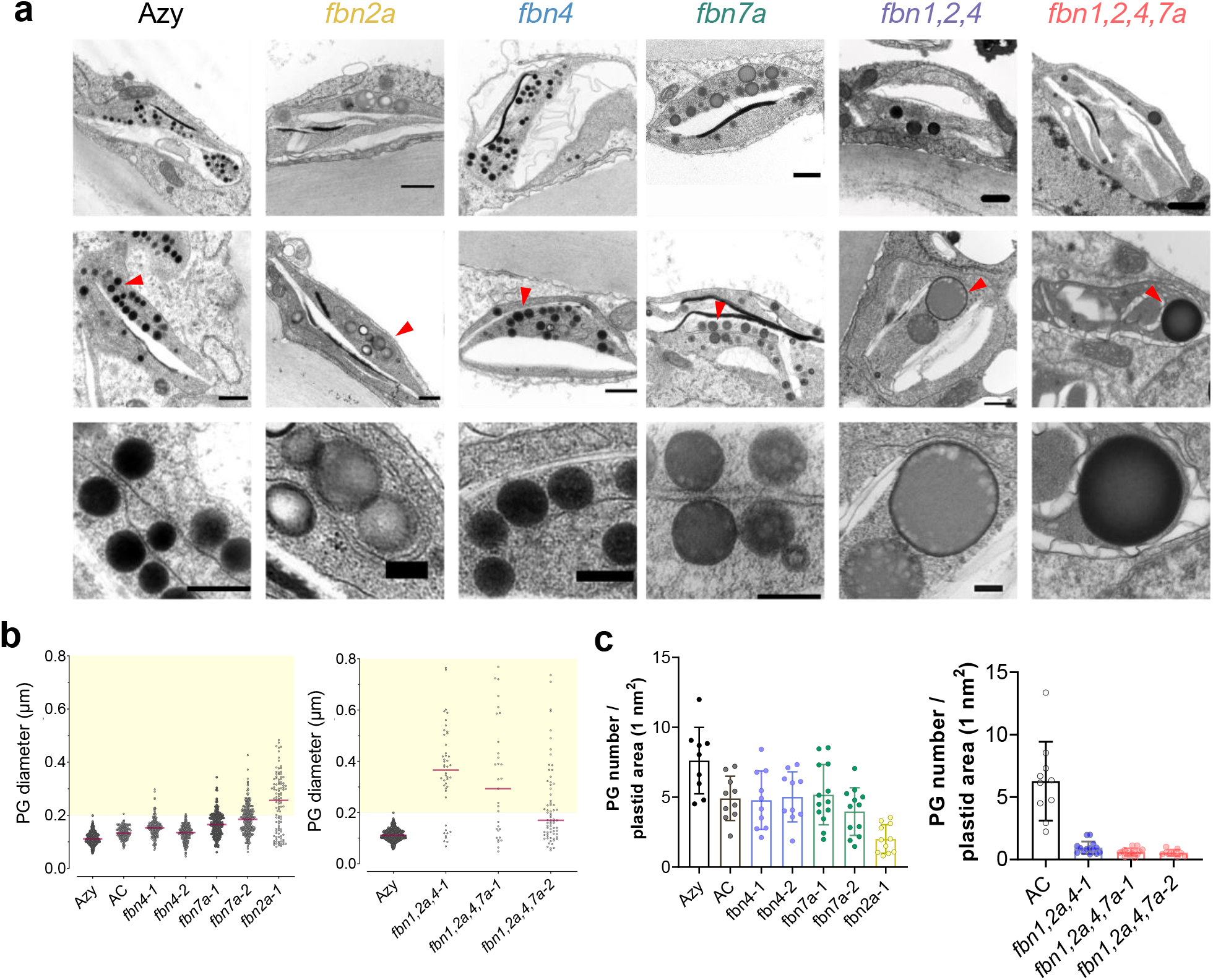
Altered fruit chromoplast ultrastructure of *fbn* mutants.

Triple and quadruple mutants displayed significant alterations in fruit chromoplast structure consistent with their carotenoid profile. They frequently failed to develop typical globular PGs. Those PGs detected were found to be unusually bigger with some even reaching giant size (diameter > 500 nm). Lycopene crystals seemed to be less developed in comparison with the control, consistent with lower lycopene levels (Fig. 6).

### Metabolome alterations associated with FBN deficiency

The impact of PG-associated FBNs on the plant metabolome focused on high-order *fbn* mutants were explored. Changes in metabolic composition of primary metabolites were addressed by GC-MS profiling of polar and nonpolar saponified extracts of fruits and leaves (SI Appendix, Table S6). In total, 117 metabolites were identified across the genotypes. Principal component analysis (PCA) using the combined dataset revealed little variation in the primary/intermediate metabolite composition of the *fbn* mutants compared to their respective controls. The exception were green fruits (MG) were a separation between *fbn* mutants and controls was attained (SI Appendix, Fig. S6). A limited number of metabolites were found to be altered in the *fbn* mutants compared to their azygous controls. In leaves, MG and ripe fruits 18, 12 and 8 metabolite differentiators were detected respectively and included representative amino acids, sugars, organic acids, isoprenoids, and fatty acids (SI Appendix, Table S6). For example, *SlFBN* deficiency led to unexpected changes in amino acid content of *fbn* fruits, these changes included increases in glutamine and threonine at the MG stage and significant decreases in phenylalanine in ripe fruits. Among the fatty acid s released through saponification, myristic acid (C14:0) decreased consistently in the *fbn* samples, while in leaves, other minor saturated species were also found altered such as pentadecanoic acid (C15:0) and heptadecanoic acid (C17:0) (SI Appendix, Fig. S6).

PGs are known to be implicated in lipid remodelling (3), therefore a deeper lipidomic analysis was performed based on LC-MS: (i) an untargeted lipidome, and (ii) a specific targeted analysis for triacylglycerol (TAG) and its derivatives.

PCA from untargeted analysis could not discriminate between the *fbn* mutants and their controls (SI Appendix, Fig. S7). Orthogonal partial least squares discriminant analysis (OPLS-DA), created a separation between the control, triple and quadruple *fbn* mutant lipidomes. The highest variable on projection (VIP) scores, which rank the importance of features in distinguishing genotypes, were retrieved. Furthermore, the correlations patterns of increasing and decreasing lipid abundance (*i.e*., Pearson’s correlation) as a function of *fbn* mutants were checked. These combined analyses revealed *SlFBN* deficiency had the strongest effects on plastidial membrane galactolipids (monogalactosyldiacylglycerol, MGDG; digalactosyldiacylglycerol; DGDG) and TAG; the latter presumably representing pools derived from the cytosol and plastids (SI Appendix, Fig. S7). In leaves, while several MGDG and DGDG related species increased in both triple and quadruple mutants, several TAG species were found depleted particularly in quadruple mutants. The same trend for the TAG lipid class was found in petals, but it contrasts to leave tissues, MGDG and DGDG related lipid species featured among the top representatives decreased in *fbn* mutants. Targeted TAG analysis (SI Appendix, Fig. S8) allowed further confirmation of impaired esterification capability associated with defective *fbn7a* alleles; several TAG-related species were found exclusively depleted in quadruple mutants either in leaves or flowers. Notably, lowered TAG-related lipid species were found both in triple and quadruple mutants, suggesting that aberrant PG morphologies associated with *SlFBN* deficiency *per se* are sufficient to trigger changes in lipid metabolism. Additionally, TAG-specific *fbn1,2a,4,7a* changes implies that FBN7a may modulate broader lipid esterification not only carotenoids.

### Expression of genes encoding PG-located enzymes are not altered in *fbn* mutants

To explore potential compensatory molecular mechanisms triggered by the targeted loss of *SlFBN*s, the expression of other *SlFBN*s encoding PG-associated proteins in flowers were compared (SI Appendix, Fig. S9). In petals, lower mRNA levels of the target *SlFBN*7a, highly expressed in flowers, were found in *fbn*7a and quadruple *fbn*1,2a,4,7a mutants. Importantly, transcript levels of *SlFBN*7b and *SlFBN*8 displayed no significant perturbation in *fbn* mutants, suggesting a lack of compensatory transcriptional response. Moreover, expression of the genes encoding PG-located biosynthetic enzymes, including *SlCCD4*B and *SlPES*1/*PYP* remained largely unchanged, except for a low level decrease of *SlPES*1 mRNA levels found in *fbn*7a mutants. This latter gene transcript encodes the acyltransferase responsible for xanthophyll esterification in tomato flowers (31). Expression of *PSY*1, the gene encoding the first enzyme of the carotenoid pathway, also remained unchanged in *fbn* mutants. These results suggested lack of major transcriptional compensatory mechanisms triggered by loss of functional *SlFBN*s.

### Proteome analysis of *fbn* mutants exposes abnormal protein targeting to PG surface

To investigate the direct changes caused by lack of functional PG-related FBNs on PGs, proteomic analysis was performed on fruit sub-plastidial chromoplast fractions, focusing on PG-enriched fractions. The composition of tomato fruit PG proteome was defined based on plastid proteins consistently identified in AC and azygous samples, with potential contaminants filtered out according to subcellular localisation data (SI Appendix, Table S7). From the SlFBN family, four members, SlFBN1, −2a, −4 and −8, were readily identified as PG-associated proteins. Surprisingly, SlFBN7a was not detected neither in the PG fraction nor further membrane-related fractions. Other PG-associated proteins (11) including NAD(P)H-ubiquinone oxidoreductase C1 (NDC1), tocopherol cyclase (VTE1), PYP/PES1, plastoglobular protein of 18 kD (PG18), and flavin-reductase related (FRR). Moreover, the vesicle-inducing protein in plastids 1 (VIPP1) was found. This protein is known for its roles in the biogenesis and repair of thylakoid membrane protein complexes and conferring tolerance to membrane stress (36), and therefore considered as a putative PG-associated protein.

Both in triple and quadruple *fbn* mutants, target SlFBN1, −2a, and −4 were absent at all analysed fractions. Indeed, only a few PG representatives were detected in *fbn* mutant proteome. This included FRR, a PG unknown for the triple and LOXB for the quadruple mutants. Thus, PG protein composition is largely controlled by mechanisms that require functional FBNs.

## DISCUSSION

The present work has exploited the CRISPR/Cas-based technology to systematically interrogate the functionality of the *FBN* multigene family. The repertoire of edited alleles generated (Fig. 2, SI Appendix, Table S3) both as individual and higher order mutants represent a valuable genetic resource and contributes to provide information about the protein structure (Fig. 3, 5).

The comprehensive characterisation of *fbn* mutants clearly indicated that FBN are structural components of the PG but functional redundancy exits. For example, multiple *SlFBN* deficiencies such as the combination of *SlFBN* 1, −2a and 4 result in heterogenous and supersized PGs in the higher-order *fbn* mutants, contrasting the normal PGs found in single mutants (Fig. 5, 6; SI Appendix, Fig. S5). Whether the *fbn* deficiencies cause dysfunctional PG biogenesis and/or increased coalescence remains to be determined. FBNs have previously been implemented in preventing PG coalescence (26), and PG formation via budding from the thylakoid membrane (37). This hypothesis is supported by the present study from the overrepresentation of dismantled thylakoids resulting in discontinuous monolayers in the irregular PGs, occurring in higher-order mutants (Fig. 5D).

The characterisation of the higher-order mutants and spatial proteomics support SlFBN1, −4, −2a and −8 as components of the core PG proteome (Fig. 4, SI Appendix, Table S7). For SlFBN7a, despite PG-association supported by the SlFBN7a-YFP fusion expression (Fig. 1), trypsin-generated peptides were not detected in our fruit proteomic data. This contrasts with a previous analysis of fruit proteome in which SlFBN7a has been identified in tomato (37). SlFBN7a absence could be explained either by lower SlFBN7a protein levels in the analysed tissue sub-chromoplast compartments and/or differences in technical detection (e.g., GEL-LC versus in-solution digestion). Notably, SlFBN7a mostly accumulated in flower chromoplasts rather than fruits based on the transcriptional profile (Fig. 1) which directly coincides with the prevalent flower defects observed in lines carrying loss-of-function *SlFBN7a* alleles. A common feature of the present spatial proteomic data and that of others is the partitioning of PG-localised SlFBNs between PG and other plastidial sub-compartments (Fig.1, SI Appendix, Table S7). The data suggests that the PG proteome can dynamically change during plastid development/environmental cues (3), raising questions about how proteins are targeted to PGs. It may rely on biophysical/chemical properties of amphipathic helix-containing proteins (38) and also influenced by post-translational modifications (39), since no dedicated protein-targeting/import machinery has been identified so far.

Notably, our data indicated that *SlFBN*7a deficiency led to dramatic loss of carotenoid ester formation in flowers. The enzyme responsible for carotenoid esterification in flowers is PES1 (31, 40), a non-specific acyltransferase that is PG localised. The question now raised is the nature of the association/interaction of *SlFBN*7a with PES. For example, is the link direct interaction modulating activity, either by facilitating an active structural conformation or precursor supply. Interestingly further reductions in carotenoid esters were observed with triple and quadruple mutants containing *SlFBN*7a. Therefore, it would appear for optimal activity a specific macromolecular configuration is favoured. This view is also supported by the finding that SlFBN7a modulation goes beyond carotenoid esterification with several TAG-related species being significantly decreased both in leaves and flowers of quadruple mutants carrying defective *fbn*7a alleles (SI Appendix, Fig. S7, S8).

In a similar fashion to the effect of SlFBN7a on carotenoid esterification, deficiencies in SlFBN2a led to reduced PC-8 levels. The AtFBN2 has been shown to interact with NDC1, which reduces the oxidised PQ-9 pool in PGs. Reduced PQ-9 is further used in PC-8 production by tocopherol cyclase (VTE1) (41, 42). Thus, SlFBN2a could positively control PC-8 formation by the direct control of NDC1 or indirectly by NDC1’s access to the plastoglobular non-photoactive PQ-9 pool. Interestingly, the functional role of AtFBN5 and the PQ-9 biosynthetic enzyme solanesyl synthase (15) interaction is still unclear. Notably, NDC1 protein was detected in membrane-enriched fractions of *fbn* mutants (SI Appendix, Table S8). In contrast to SlFBN2a, the absence of SlFBN7a resulted in increased PC-8 levels. Potentially, this finding could arise from a re-direction of precursors into prenyl lipids because of reduced carotenoid esterification following the absence of SlFBN7a. Alternatively, the increased PC-8 in SlFBN7a defective lines could be a compensatory effect derived from altered TAG composition. For many years sectors of isoprenoid biosynthesis have been associated with biosynthetic metabolons and a requirement for molecular scaffolds (reviewed in (43)). Evidence from SlFBN2a (and AtFBN2, (42)), SlFBN7a and AtFBN5 (15), as well as the recently identified agal chloroplast SEC14-like1 protein (CPSFL1) (44) and tomato tocopherol binding portein (SlTBP) (45) now opens the potential for a range of biosynthetic ancillary proteins, that could bind and/or transfer substrates directly to enzymes directly or indirectly as part of large macromolecular structures.

Cellular ultrastructure analysis revealed dramatic effects upon PG resulting from the absence of multiple FBNs. However, very few perturbations to the metabolome arose. The most pronounced changes occurring in the lipidome, which is not surprising considering the nature and postulated function of PGs. The depletion of myristic acid in *fbn* mutants (SI Appendix, Fig. S6), one of the fatty acid moieties incorporated into TAG and phytyl esters presumably via intermediates derived from fatty acid *de novo* synthesis (46), was a first indication that FBN deficiency led to alteration in lipid metabolism potentially due to lack of PG lipid partitioning. Most strikingly, are the common alterations in galactolipids and TAG species shared by triple and quadruple mutants (SI Appendix, Fig. S7, S8). In the former case, a tissue-specific response was detected, with MGDG and DGDG species reduced in *fbn* mutant flowers but increased in leaves. In both cases plastidial galactolipid turnover into TAG (46, 47) may be compromised given alterations in PG structure and associated processes. It is possible, however, that flower and leaf differences in galactolipids are caused by differential metabolic demands between these plastid types. It is interesting that in many metabolic engineering outputs, targeting isoprenoids (carotenoids) the resulting changes in metabolite composition are accompanied by dramatic plastidial ultrastructural perturbations and altered transcript levels of key pathway genes (22, 48). Here, under non-stressed conditions, dramatic changes in sub-plastidial structures have had relatively minor holistic effects to the steady state metabolome or key related transcript levels.

In summary, multiplex gene-editing tools have been used to decipher the functionality of the FBN multigene family. Functional redundancy is evident, but collectively FBNs clearly play a major role in PG biogenesis and formation. In addition, the data implements specific SlFBNs (*e.g*. −*2a* and −*7a*) in carotenoid esterification, TAG utilisation and PC-8 formation. The overall cellular and metabolic effects of the FBNs tested are summarised in Fig. 7. Our findings extend knowledge on fundamental aspects of metabolic compartmentalisation in plant cells revealing the importance of lipoprotein particles for plastid metabolism/physiology. The genetic resources generated from this present work paved the way to elucidate to role of FBNs and the PG, in combating biotic and abiotic stresses as well as future strategies for the development of plant cell factories producing nutritional and industrially valuable compounds.

**Fig 7.**
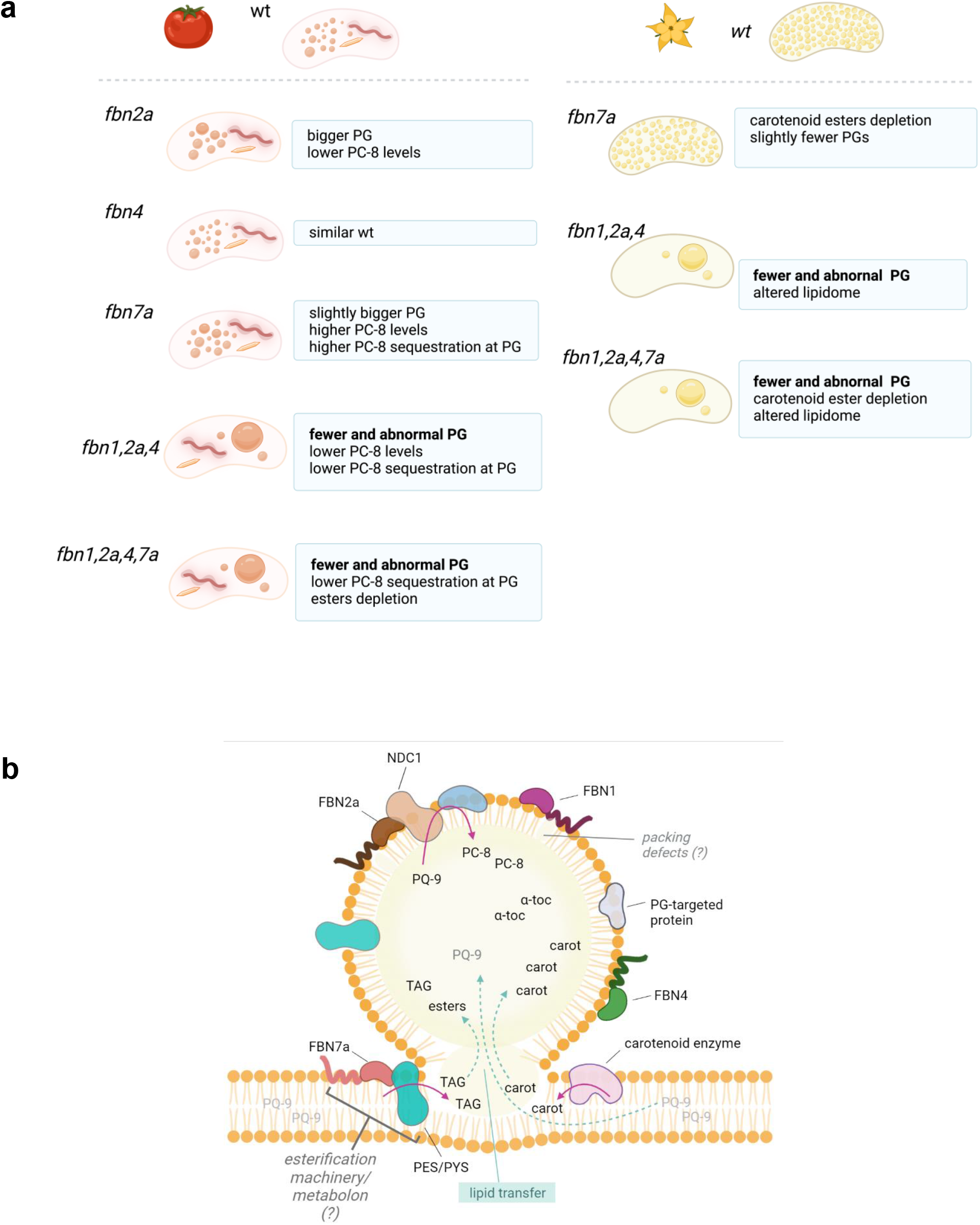
Summary of changes associated with SlFBN deficiency and proposed model.

## METHODS

### Phylogenetic analysis

For phylogenetic analysis, BlastP searches were performed using the protein sequences of *Arabidopsis thaliana* FBN (3) as queries against the tomato genome (Solanaceae Genomic network, http://solgenomics.net). For pepper (*Capsicum annuum*, cv. Zunla), sequences were retrieved from PepperHub database (27). The sequences were aligned using the MUSCLE package available in the MEGA X software with default parameters, and Neighbour–Joining phylogenies with 1000 bootstrap replications were created with the distances calculated according to the best model indicated by MEGA (49).

### Subcellular Localization and confocal microscopy

For SlFBN subcellular localization, the full-length CDS of SlFBN2a, SlFBN4, SlFBN7a (without stop codon), were cloned into the p2YGW7 resulting in a C-terminally tagged YFP protein. The PG marker, AtFBN1a fused in-frame with CFP, used was from the vector p2CGW7 (28). Arabidopsis mesophyll protoplasts were isolated as previously described (50). Protoplast transfection assay was performed using a polyethylene glycol-based method (51) immediately after the protoplast isolation. After 24h, fluorescence emitted from the protoplasts was observed using a TCS-SP8 laser scanning confocal microscope (Leica Microsystems CMS, Germany) as follows: excitation 514□nm, emission 520–570□nm for YFP; excitation 458□nm, emission 465–505□nm for CFP; excitation 514□nm, emission 650–740□nm for chlorophyll autofluorescence. Emitted fluorescence was false□coloured in green (YFP), red (CFP) and blue (chlorophyll). The confocal images shown are maximal projections of selected planes of a *z*□stack.

### Plant material and growth conditions

Tomato plants (*Solanum lycopersicum*, cv. Ailsa Craig) were grown under greenhouse conditions with a 16/8-h day night photoperiod at 25°C /19°C, respectively. For experiments, homozygous/biallelic and azygous segregants were grown at the same time in the same glasshouse room.

### CRISPR-Cas9 constructs for generating tomato fbn mutants

Genomic sequences 5’-G(N)_19_NGG were selected to design two gRNA spacer sequence targeting specifically SlFBN1, SlFBN2a, SlFBN4, SlFBN7a, avoiding off-target effects, using CRISPR-P software (52). Secondary structure analysis of target-sgRNA sequences was performed with the program RNA folding tool (53). The standardised modular cloning system Golden Gate MoClo was used (54, 55). The Cas9 expression cassette consists of a 2×35S Cauliflower Mosaic Virus promoter (pICH51288) and a plant codon-optimised Cas9 coding sequence, pcoCas9 (56), which was domesticated for removal of internal BbsI and BsaI restriction sites, both parts used to assemble a Level 1 transcriptional unit (pICH47742::2×35S::Cas9) according to (57). For the gRNAs, a 5’ tailed primer harbouring the individual *SlFBN* spacer (guide sequence), BsaI recognition site and compatible vector overhang was used to generate specific gRNA amplicons by PCR from an gRNA scaffold template (AddGene no. 46966; for primers, see SI Appendix, Table S9). Each gRNA PCR fragment was individually assembled with Arabidopsis U6 small RNA promoter (pICSL01009::AtU6p, AddGene no. 46968) in the appropriate Level 1 vector (plCH47751, plCH47761, plCH47772, plCH47781, pICH47791, pICH47732, pICH47742, pICH47751) depending on the position of each gRNA in the final binary vector as described by (57). A neomycin phosphotransferase II (NPTII) cassette (pICH47732::NOSp::NPTII, Addgene no. 51144) was used as selectable marker for plant transformation. For single *fbn* mutants, NOSp::NPTII; 2×35Sp:Cas9; Level 1 AtU6p::gRNA1, Level 1 AtU6p::gRNA2 were assembled into the final Level 2-based binary vector. For building multiplex *SlFBN* gRNA vectors, NOSp::NPTII; 2×35Sp:Cas9; level 1 AtU6p::gRNA1 to AtU6p::gRNA(n) were assembled into Level P vector, and produced multiplex vectors P1-M1 (designed for targeting *SlFBN*1, *SlFBN*2a, *SlFBN*4) and P1-M2 (designed for for targeting *SlFBN*1, SlFBN2a, SlFBN4, SlFBN7a). Restriction-ligation reactions were performed in 20 μL volume in a thermocycler for 40 cycles of 37 °C for 3 min followed by 16 °C for 4 min, with a final incubation of 5 min at 50 °C and 5 min at 80 °C. All assembled plasmids were validated by restriction enzyme analysis and confirmed by Sanger sequencing. Final binary vectors were transformed into *Agrobacterium tumefaciens* (strain LBA4404).

### *Agrobacterium tumefaciens*-mediated transformation and selection of transformants

*A. tumefaciens*-mediated transformations of the *S. lycopersicum* cv Ailsa Craig were performed as previously described in (58) using cotyledon segments from 8-day-old seedlings precultured with Agrobacterium containing the CRISPR/Cas9 constructs of interest. Selective regeneration medium containing kanamycin (100 mg/L) was used for explant selection. After 8 weeks, well-rooted T_0_ plantlets were acclimatized to greenhouse conditions and their leaves sampled for genotyping.

### Detection and determination of targeted mutagenesis

Genomic DNA was purified from leaves using the DNeasy 96 plant kit (Qiagen) following manufacturer’s instructions. The presence/absence of the transgene in regenerated primary plants (T_0_) and offspring (T_1_ and T_2_ plants) was detected by PCR in the genomic DNA using NPTII and Cas9□specific primers (SI Appendix, Table S9). To characterize the Cas9□induced mutations in the transformed plantlets, specific primers for *SlFBN*, whose binding positions are about 250 bp upstream of the target site, were used for PCR amplification. Amplicons were analysed by Sanger sequencing. For those with multiple peaks in chromatograms, the PCR product was cloned into TOPO-TA (ThermoFisher Scientific) and, at least five clones per sample were sequenced. T_1_ progeny of the confirmed T_0_ gene-edited lines was PCR-genotyped for screening of inherited mutant alleles (either in homozygous or biallelic state) and Cas-9 null segregants. Azygous segregant line descended from an original CRISPR-single and from a CRIPSR-multiplex vector transformant were selected as control lines. Phenotyping was performed on T_2_ Cas-9 null segregant homozygous offspring for *fbn* singles, triple and quadruple mutants, unless stated otherwise. For multiplex CRISPR constructs, an extra set of gene-edited lines for triple *fbn*1,2a,4), quadruple (*fbn*1,2a,4,7a) as well as double mutant (*fbn2a,4*), the latter generated through a P1-M1 transformants whose SlFBN1 edition failed, was obtained in Ailsa Craig background lacking green shoulder (glk2/uniform recessive). Their corresponding T_1_ progeny harbours Cas9-gRNA expression cassette Cas^+^.

### Isoprenoid determination and quantification

Isopre*n*oids (carotenoids, tocochromanols, and chlorophylls) were extracted from lyophilised tissue powder (15 mg) as described by (23). For ultra-performance liquid chromatography (UPLC)-Photo Diode Array (PDA) method, compounds were analysed by reverse-phase chromatography using an UPLC system (Acquity, Waters) equipped with PDA detector (Acquity, Waters). A UPLC BEH-C18 column (100 mm × 2.1 mm; 1.7 μm, Acquity, Waters) was used for separation as described in (24). For high performance liquid chromatography (HPLC) method, analyses were conducted as described in (58). Peak identification was achieved by comparison of characteristic UV/Vis spectrum with authentic standards, reference spectra and retention times (48). For xanthophyll esters in flowers, a spectra table is available in SI Appendix, Table S10. Quantification was performed using dose-response curves obtained from authentic standards when available.

### Subchromoplast fractionation

Chromoplasts were isolated from fruits (90 g) at B3 to B4 stage, and subcompartments were fractionated using a discontinuous gradient of sucrose, according to (24).

### qPCR expression analyses

Total RNA was extracted from flowers and fruit pericarps using the RNeasy kit (Qiagen) according to manufacturer’s instructions. RNA quality was assessed by agarose gel electrophoresis. Total RNA (1 μg) was treated with DNase and converted into cDNA using the QuantiTect Reverse Transcription kit (Qiagen), according to the manufacturer’s protocols. Real-time quantitative PCR (qPCR) assays were performed in duplicate using RotorGene SYBR green PCR kit (Qiagen) on Rotor-Gene Q, with approximately 10 ng of reverse-transcribed RNA. Primer sequences are listed in SI Appendix, Table S1. Relative expression was calculated as described by (59). For reference gene selection, expression stability of four known reference genes (CAC, EXP, ACT1 and ACT2) (60, 61) were evaluated on control and heat-stressed samples using GeNorm (62). ACT2 and CAC were selected based on lowest expression stability values (M) of 0.362 and 0.438, respectively.

### Transmission Electron Microscopy

Segments from pericarp fruit and mature, opened flowers were fixed at room temperature in solution [3% (v/v) glutaraldehyde, 4% (v/v) formaldehyde buffered with 0.1 M PIPES buffer pH 7.2] and then stored at 4 °C for at least 24 h until processing. Samples were post-fixed in buffered 1% (w/v) osmium tetroxide and uranyl acetate, washed, dehydrated in a graded series of acetone, and embedded in resin. Ultrathin sections were stained with Reynolds lead citrate and imaged on a Tecnai T12 Transmission Electron Microscope (Field Electron and Ion Company, USA). Measurements were made in ImageJ v.2.0.0-rc-68/1.52e (National Institutes of Health).

### Immunoblotting and Proteomic analysis

For immunoblotting, ripe fruit homogenates (150 mg) were extracted with extraction buffer containing 0.1 M Tris-Cl (pH 7.5 − 8.0), 6 M urea, 2 M thiourea, 0.2% (v/v) Triton X-100, 0.2% (w/v) sarcosyl and 2 mM DTT. Protein was quantified using the Bradford method (63), and 10 μg were separated on a 12.5% polyacrylamide gel and blotted to PVDF membrane. Blots were probed for the presence of the FBN1 using a polyclonal antibody raised against *Capsicum* FBN1/PAP as described by (64).

For proteomic analysis, total protein was extracted from isolated fractions, run in the SDS-PAGE gel, subsequently digested with trypsin and analysed into nano LC-MS/MS as described by (24). Analyses were conducted in an AdvanceBio Peptide Map column (2.1×100mm, 2.7micron, Agilent Technologies, Inc.) and Infinity II 1290 UHPLC coupled to a 6550 iFunnel QTof (Agilent Technologies, Inc.). Data analyses were performed using Spectrum Mill MS Proteomics Workbench (Rev B.06.00.201, Agilent Technologies, Inc.) and Mascot Distiller (v2.4.2.0, Matrix Science) as described in (24).

### Metabolite profiling by gas chromatography GC-MS

Polar and nonpolar saponified extracts from fruits and leaves were prepared and analysed as previously described in (65).

### Lipid analysis by liquid chromatography (LC)-MS/MS

Leaf, fruit and flower petals (5-10 mg) were extracted with chloroform: methanol (1mL, 2:1, v/v). Analysis was performed as previously described by (66) for phospho- and galactosyl lipids and neutral lipids. Samples were processed with Agilent Profinder (v10.0 SP1, Agilent Technologies, Inc.) and identification performed through comparison to authentic standards and MS/MS fragmentation pattern.

### Statistical analyses

Univariate statistical analysis for comparison between control and *fbn* mutants was performed by Student’s t test or ANOVA followed by a posthoc test (as stated in figure and tables) with the level of significance set to 0.05, using GraphPad Prism 8 software. For multivariate statistical analysis, MetaboAnalyst 4.0 (67) and Simca®17 (Sartorius) softwares were utilised.

## Supporting information

Supplemental data sets

## Data availability

Raw data files for lipidomics and proteomics are available through Mendeley data (DOI: 10.17632/9xmnybzgnd.1 and DOI: 10.17632/3hxz7c77r5.1). All other data generated or analysed during this study are included in this article (SI Appendix).

## Acknowledgements

This work is supported by the H2020 programme TomGEM / 679766; A holistic multi-actor approach towards the design of new tomato varieties and management practices to improve yield and quality in the face of climate change, and Biotechnology and Biological Sciences Research Council (BBSRC) OPTICAR Project (optimisation of tomato fruit carotenoid content for nutritional improvement and industrial exploitation; Project BB/P001742/1). Drs Genny Enfissi and Marilise Nogueira (RHUL) are thanked for their constructive advice. Dr Francesca Robertson is thanked for initial proteomic analysis and Mr Chris Gerrish for expert technical assistance. Dr Marcel Kuntz, Grenoble University for the initial antibody to FBN. pICSL4723-P1 and pAGM472 were kindly provided by Mark Youles, Sainsbury Laboratory, UK.

## Author contributions

Conceptualisation (JA & PDF), methodology (JA, LP, MD & PDF), formal analysis (JA), investigation (JA), resources (PDF), data curation (JA), writing-original draft (JA), Writing-review and editing (JA, LP, MD & PDF), visualisation (JA), project administration (PDF), and funding acquisition (PDF).

