## Supplemental figures for "The *FIBRILLIN* multigene family in tomato, their roles in plastoglobuli structure and metabolism"

### **Supporting Information for**
### The FIBRILLIN multigene family in tomato, their roles in plastoglobuli structure and metabolism


Juliana Almeida1†, Laura Perez-Fons1, Margit Drapal1, Kit Liew1¶ and Paul D. Fraser1*

[1]Department of Biological Sciences, Royal Holloway University of London, Egham, Surrey TW20 0EX, United Kingdom.

[†] Present address, CTC Sugarcane Technology Center, Fazenda Santo Antônio, S/N, 13400-970, Caixa Postal 162, Piracicaba, São Paulo, Brazil.

[¶]Present address, Division of Biological and Environmental Sciences and Engineering, 4700 King Abdullah University of Science and Technology, Thuwal 23955-6900 - Kingdom of Saudi Arabia.

For correspondence: Paul D. Fraser



**This PDF file includes:**

Fig.s S1 to S9
Legends for Datasets S1 to S10

**Other supporting materials for this manuscript include the following:**

Dataset with Tables S1 to S10

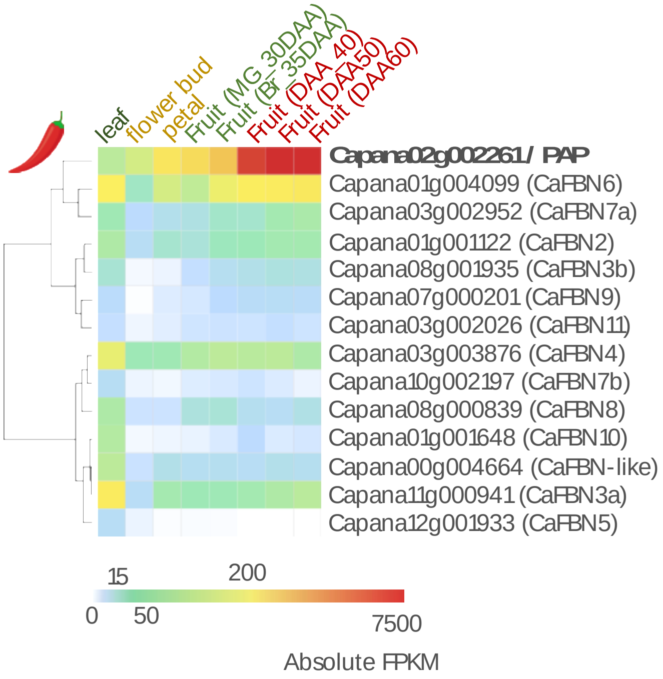


**Fig. S1. Expression profile of FIBRILLIN (FBN) loci in pepper**. Heatmap representation of CaFBN expression profile in pepper. Expression data were obtained from PepperHub (http://www.hnivr.org/), represented as FPKM-normalized values.

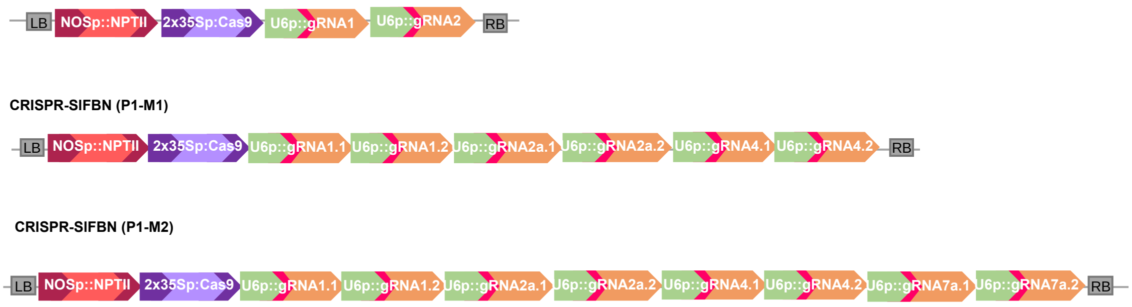


**Fig. S2. CRISPR/Cas9 constructs for target gene editing of *SlFBN* genes.** Scheme of final binary vectors containing the NPTII selectable marker, Cas9 driven by the 2x35S promoter and transcriptional units for gRNA expression (under control of U6 promoter). LB = left border; RB = right border.

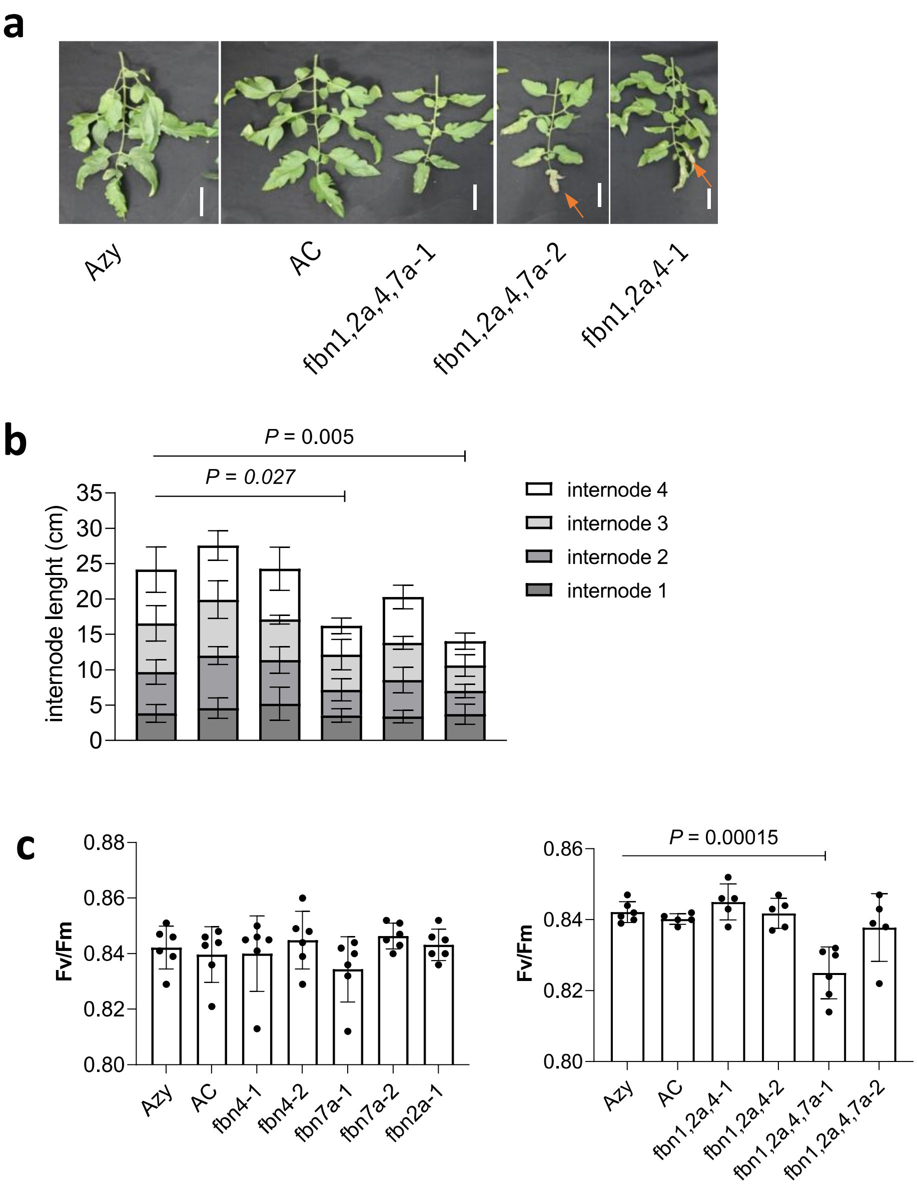


**Fig. S3. Plant morphology of multiple knockout *fbn* mutants. a** Representative fully expanded leaves (cv. Ailsa Craig) of 10-weeks-old Azy, AC and triple and quadruple *fbn* mutants. All plants were cultivated together in the same growth chamber. Orange arrows indicate lesions caused by fungal pathogens. Bar = 5 cm. **b** Internode length of the four equivalent internodes. **c** Maximum quantum efficiency of Photosystem II (Fv/Fm). Data (n > 5, biological replicates) are means and error bars indicate standard deviation (SD). Statistically significant differences between Azy and *fbn* mutant are indicate by *P*-values (one-way ANOVA with Dunnett's post-tests, $P < 0.05$).

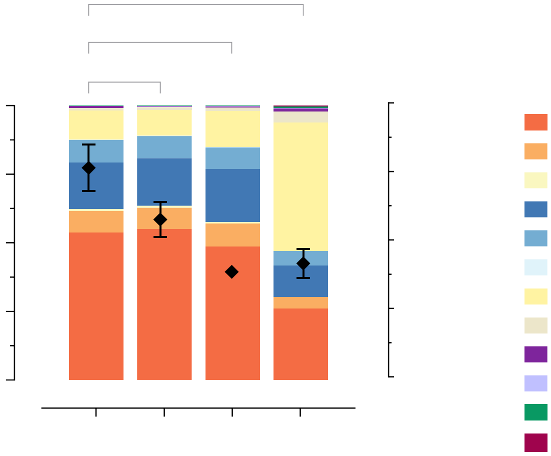

**Fig. S4. Flower carotenoid composition of *Cas*⁺/*SlFBN*-edited lines.** T1 SlFBN-edited lines profiling by HPLC-PDA. Carotenoid composition and total levels of double (*Cas+/fbn2a,4*), triple (*Cas+/fbn1,2a,4*) and quadruple (*Cas+/fbn1,2a,4,7a*) *fbn* mutants. Data (n = 3, biological replicates) were analysed by one-way ANOVA with Dunnett's post-test; asterisks denote statistically significant differences (* P < 0.05, ** P < 0.01, *** P < 0.001) compared to AC control. Error bars indicate ± SD. M, mono-esters; D, di-esters; F, free forms.

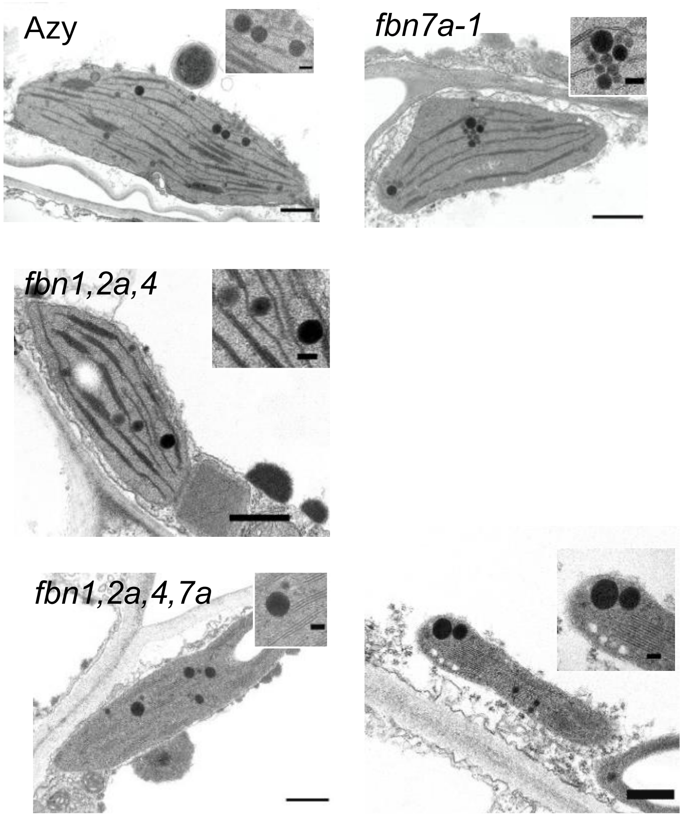


**Fig. S5. Aberrant PG morphology found in flower plastids of high-order *fbn* mutants.** Representative transmission electron (TEM) micrographs of early-stage plastids (bearing thylakoids organised in *grana* and a few electron dense PGs) from petals from Azy and *fbn* mutants. Detailed view of PGs was shown on the inset panels. Giant PG formation was observed in some plastids of quadruple (*fbn1,2a,4,7a*) mutants. Scale bars = 500 nm (main panels), and 50 nm (inset).

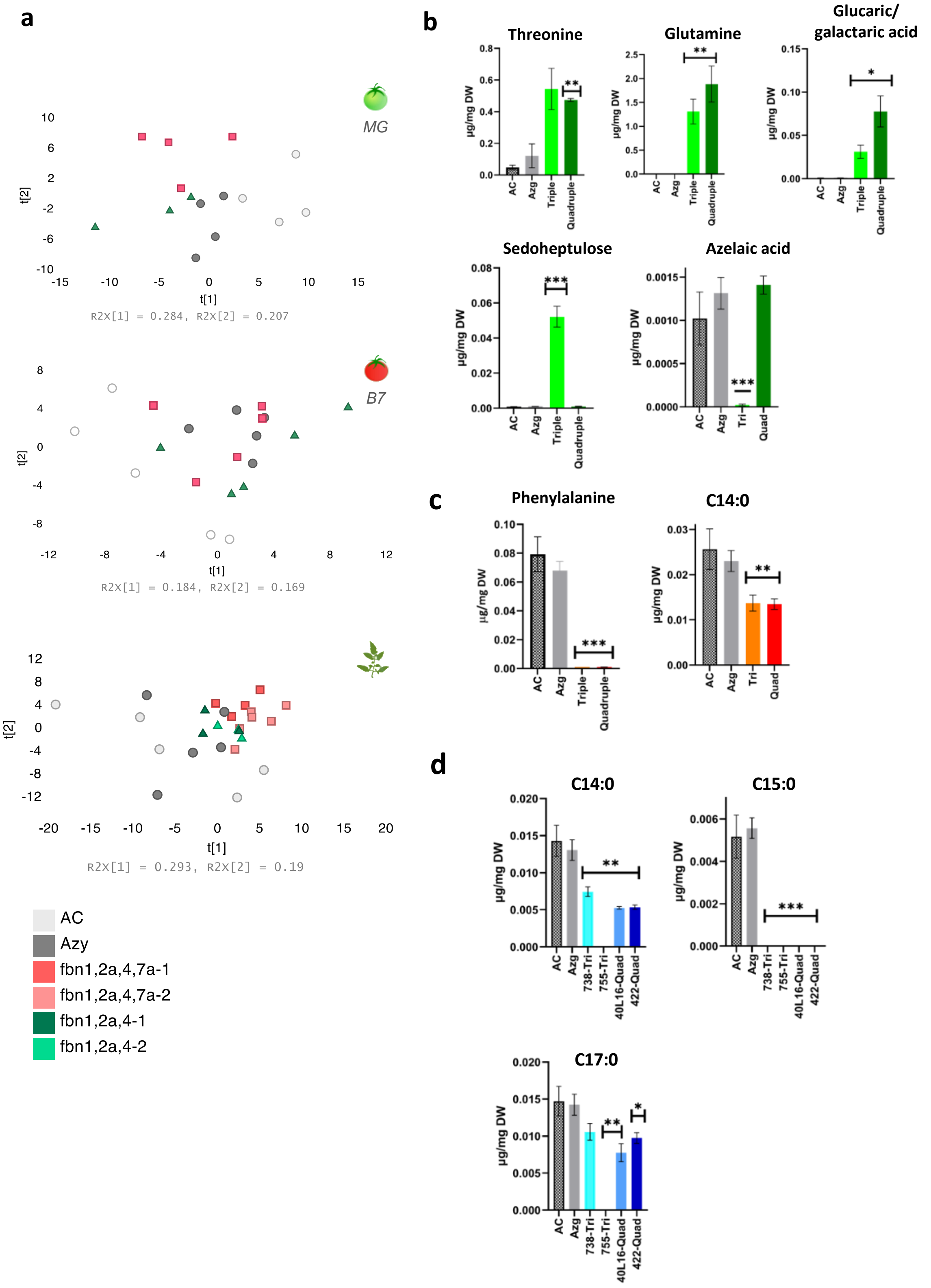


**Fig. S6. Effect of SlFBNs deficiency on tomato primary metabolism. A** Scores plot obtained by principal component analysis (PCA) for metabolite levels measured in polar and nonpolar extracts in fruits at mature green (MG) and ripe (B7) stage, and in leaves (from top to bottom). Changes in metabolites associated with SlFBNs deficiency in MG **B**, ripe fruits **C** and leaves **D**. Quantification was determined relative to the internal standard and values are presented as mean ± SD from four biological replicates. Only significant changes compared to respective Azy control are shown (pair-wise t test corrected for multiple comparison using Holm-Sidak's post-test; * Adjusted $p < .05$, ** $p < .01$, *** $p < .001$). Full data set available in Table S6. C14:0, myristic acid; C15:0, pentadecanoic acid; C17:0, heptadecanoic acid.

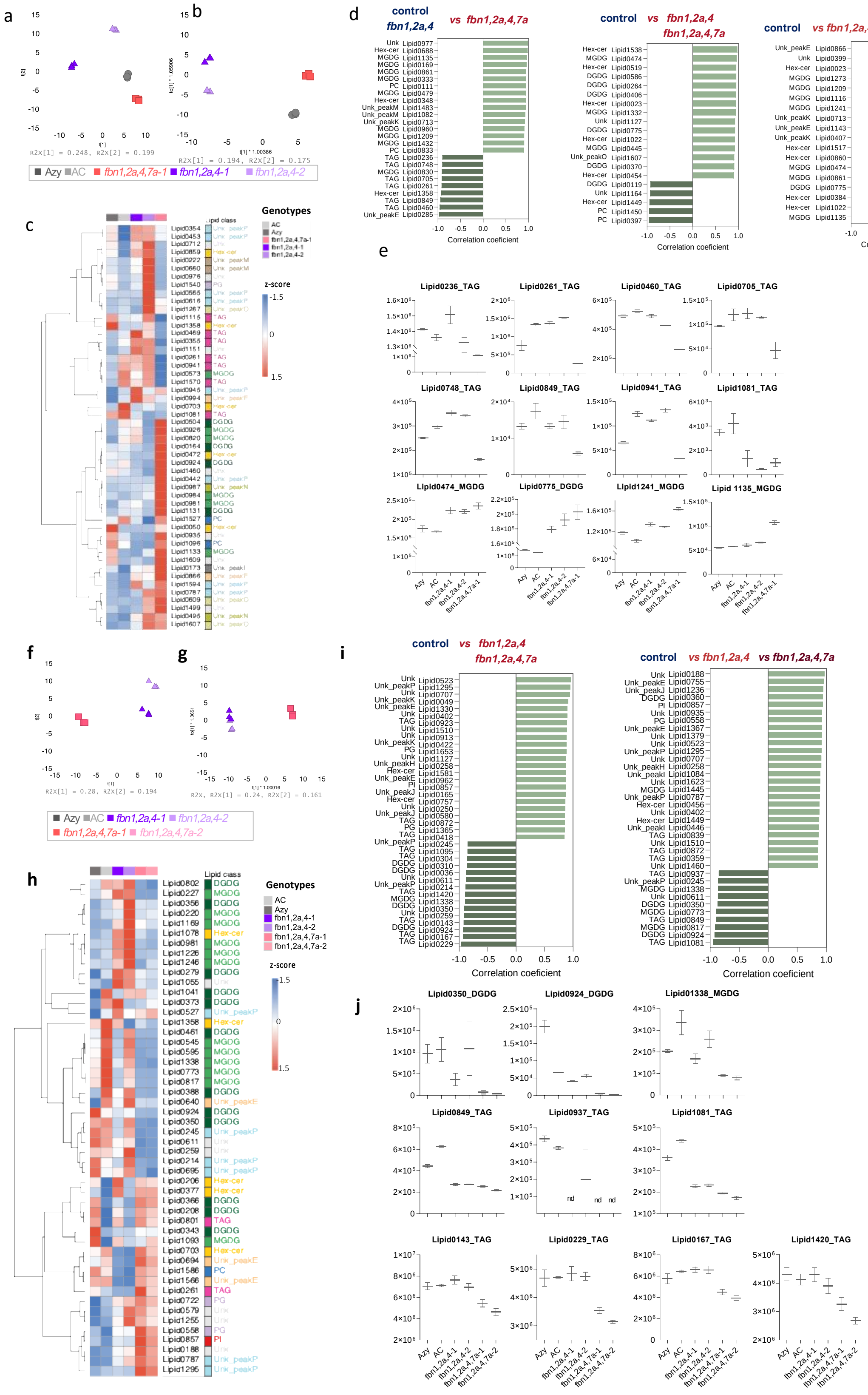

a
b
c
d
e
f
g
h
i
j
Genotypes
AC
Azy
fbn1,2a,4,7a-1
fbn1,2a,4-1
fbn1,2a,4-2
fbn1,2a,4,7a-2
z-score
Lipid class
control
vs fbn1,2a,4
vs fbn1,2a,4,7a
Correlation coeficient

**Fig. S7. Untargeted lipidomic analysis of high-order *fbn* mutants.** Data was obtained from based on hydrophilic interaction liquid chromatography (HILIC) method. **a** PCA and **b** OPLS-DA based on 1574 LC-MS chemical features found in leaves. Data were log-transformed and Pareto scaled. **c** Hierarchical clustering of top 50 variables with the highest VIP scores identified by OPLS-DA. Heat maps are coloured by red and blue colours indicating higher and lower abundance, respectively, as measured by row standardised z-scores derived from LC-MS peak intensity (n= 3 biological replicates). Genotypes are annotated as colored boxes on the top of the heatmap. The boxes on the right of the heatmap indicate the lipid classes. The red and blue colours indicate higher and lower mean expression, respectively, as measured by row standardized z-scores. **d** Chemical features that most correlate to changes that discriminate *fbn* mutants (*fbn1,2a,4* and *fbn1,2a,4,7a*) and controls (Azy and AC). Values correspond to Pearson correlation. **e** The graphs represent some changes observed on selected chemical features based on correlation or VIP scores analyses that are significantly different (ANOVA followed by Fisher's LSD, α = 0.05).

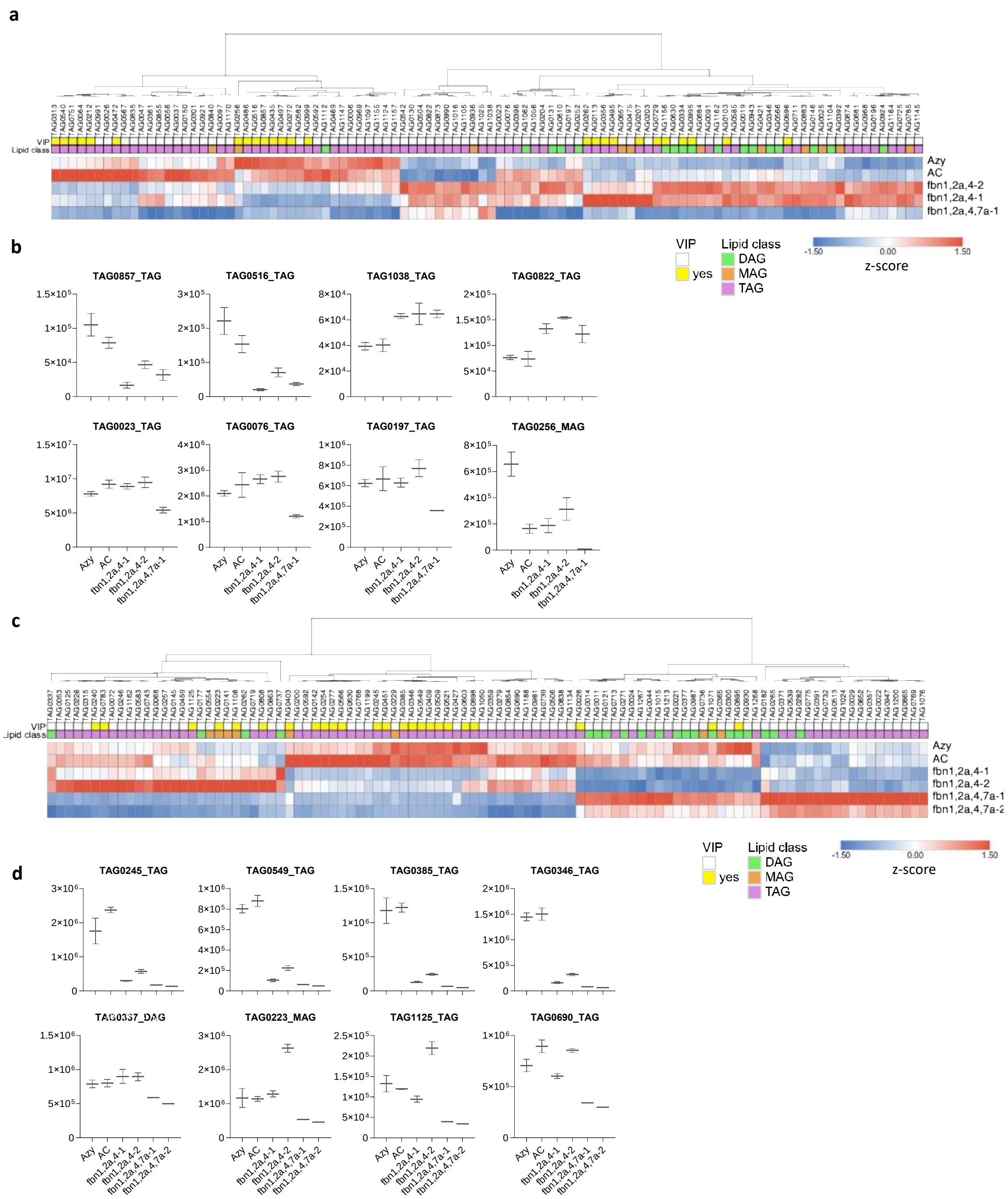

**Fig. S8. Targeted lipidomic analysis of triacylglycerol (TAG) and its derivatives. a,c** Heat map of top 100 features identified by ANOVA (Posthoc Fisher's LSD, α = 0.05, corrected by multiple comparisons) in leaf and flower, respectively. Red and blue colours indicate higher and lower abundance, respectively, as measured by column standardised z-scores derived from LC-MS peak intensity (n= 3 biological replicates). Genotypes are indicated on the right of the heatmap. The boxes on the top of the heatmap indicate the lipid classes, and whether the variable is found among the highest VIP scores (top 100) identified by OPLS-DA. **b,d** Selected chemical features from heat maps of leaf and petal, respectively, that show statistically significant changes on *fbn* mutants (ANOVA followed by Fisher's LSD, α = 0.05)

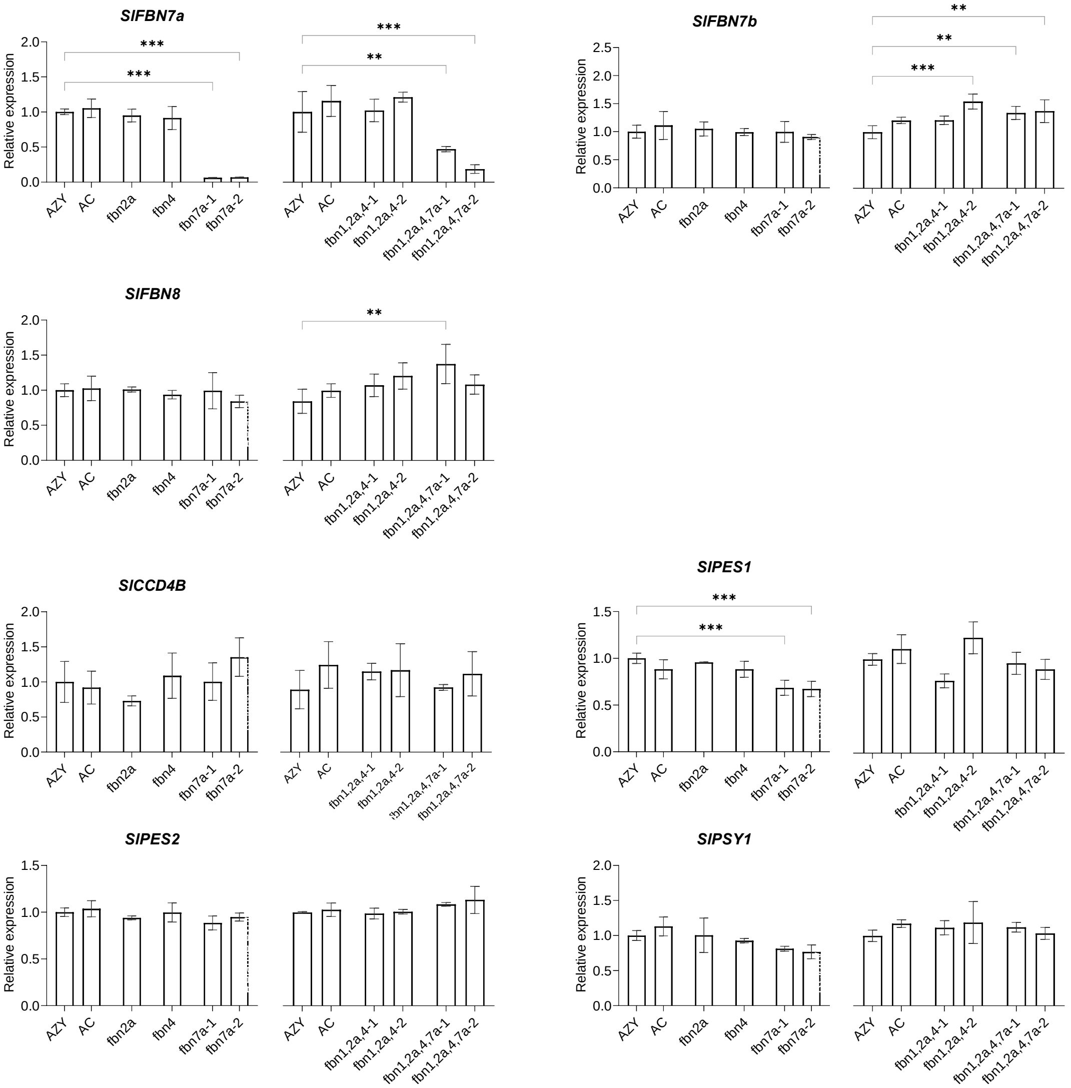
SlFBN7a
SlFBN7b
SlFBN8
SlCCD4B
SlPES1
SlPES2
SlPSY1
Relative expression
***
**
AZY
AC
fbn2a
fbn4
fbn7a-1
fbn7a-2
fbn1,2a,4-1
fbn1,2a,4-2
fbn1,2a,4,7a-1
fbn1,2a,4,7a-2

**Fig. S9. Relative expression of *SlFBNs* and other genes encoding metabolic enzymes associated to PG by qPCR.** Values are expression levels normalized to *CAC* and *ACT2* reference genes (mean ± SEM; n ≥ 3 biological replicates). Significant differences (Student's t test, * $p < .05$, ** $p < .01$, *** $p < .001$) between *fbn* mutants and control are indicated.

**Dataset S1-S10 (separated file)**

**Table S1.** gRNA recognition sequences in *SlFBN* genes and off-targets.
**Table S2.** Number of plants recovered from *Agrobacterium*-mediated transformation harbouring mutations in the *SlFBN* target gene.
**Table S3.** SlFBN edited lines phenotyped in this study.
**Table S4.** Isoprenoid profile of fruits and leaf from fbn mutants determined by UPLC-PDA.
**Table S5**. Isoprenoid profile of petals from *fbn* mutants determined by HPLC-PDA.
**Table S6**. Metabolite levels measured in fruits (MG and B7 stage) and leaves by GC-MS.
**Table S7**. Proteome associated to PG fraction of tomato fruits from control and *fbn* mutant genotypes.
**Table S8**. Proteome associated to membrane fraction of tomato fruits from control and *fbn* mutant genotypes.
**Table S9**. Primers used in this study.
**Table S10**. Peak identification of isoprenoids analysed by HPLC-PDA.
